# Introns Control Stochasticity in Metazoan Gene Expression

**DOI:** 10.1101/746263

**Authors:** Bryan Sands, Soo Yun, Alexander R. Mendenhall

## Abstract

Introns are noncoding DNA elements found in between coding exons of most genes in most metazoans. Introns are known to both increase expression levels and provide means for making different protein products from the same gene via alternative splicing. It is thought that evolution has selected for these functions of introns. Here we present evidence for an additional reason why so many metazoan genes have introns, and why some do not. We found that introns control stochasticity in metazoan gene expression. They do this by controlling stochastic autosomal allele bias. We determined this by measuring the expression of differently colored reporter alleles with and without introns. We find that introns control stochastic autosomal allele biases using a 5’-position dependent mechanism. Intron control of allele expression noise provides an additional mechanism to consider when pondering why so many metazoan genes have introns and why some have lost them.

## INTRODUCTION

Many discrete traits are incompletely penetrant, and this includes many human diseases. For example, dominant negative oncogenic mutations associated with Noonan’s syndrome and esophageal cancer are not 100% penetrant^1,2^. Furthermore, differences in genes and environments do not account for differences in complex traits, including lifespan^3,4^ and complex human diseases^5^, such as Parkinson’s disease^6^. That is, when genes and environments are held as constants in laboratories, organisms still develop differences in both complex and discrete traits. Underlying this are differences in the expression of genes (e.g., penetrance of *skn-1* mutations in *C. elegans*^7^). Consequently, there has been a significant amount of experimentation dedicated to defining and understanding the intrinsic, natural axes and mechanisms of variation in gene expression. Systems biology studies in *E. coli, S. cerevisiae* and *C. elegans* have dissected the physiological mechanisms of cell-to-cell variation in gene expression into experimentally tractable bins^8-11^.

One of the major mechanisms by which gene expression can vary in eukaryotic cells is stochastic autosomal allele expression bias, which we and others refer to and quantify as intrinsic noise^8^. When intrinsic noise is high for a particular gene, cells within a tissue will present a spectrum of gene expression possibilities, ranging from fully monoallelic (single allele expressed while the other is silenced) to balanced biallelic expression (1:1 ratio of alleles). When intrinsic noise is low for a particular gene, the majority of cells within a tissue will present as biallelic. This intrinsic noise is distinct from parental imprinting or X inactivation, and is reviewed from different perspectives here^12^, here^13^ and here^14,15^. These different groups refer to this phenomenon, or aspects of it, as intrinsic noise^12^, stochastic autosomal allele expression bias^13^, allele specific expression^14,15^, or monoallelic expression^12-15^.

Intrinsic noise is a quantitative measure of the deviation from a perfect 1:1 ratio, visible as deviation from the diagonal trend line in a scatter plot of allele expression values. This is demonstrated in images and scatter plots from single cells in^8^. When Raser and O’Shea quantified intrinsic noise in yeast, they found that alleles controlled by the PHO5 promoter were expressed along a continuum, ranging from monoallelic to biallelic. Later, genome wide SNP arrays, RNA-seq and ChIP-seq surveys of human and murine cells found extreme stochastic allele bias (often monoallelic) to be widespread^16-21^. This extreme allele bias can have many consequences, including: loss of allele-specific biochemical function, altered biochemical capacity conferred by different dosages of gene product, escape from a dominant negative phenotype, or condemnation to the manifestation of a negative recessive phenotype. Quantifying the baseline of allele bias and the factors that affect it are critical for understanding missing heritability and incomplete penetrance of human diseases.

Introns are noncoding DNA elements found between most coding regions of a gene. Work from us^22^ and others (reviewed in^23,24^), has shown that, in addition to the role introns have in alternative splicing, introns act to increase expression levels. Moreover, the presence of short introns close to the transcription start site (TSS) are associated with active chromatin marks, increased transcription and more accurate TSS usage^25^. Given these particular functions of introns in gene expression control, we hypothesized that introns could also affect the probability that both alleles of a gene are expressed. If true, then introns should decrease intrinsic noise. To test this hypothesis, we took the approach of quantifying intrinsic noise at the protein level, for reporter allele pairs with and without introns, in the tissues of live *C. elegans*.

In this study, we found intrinsic noise to be visually detectable in microscopic images of cells in *C. elegans* tissues. We found that introns can decrease intrinsic noise by an order of magnitude. We found this to be true for synthetic and natural introns. We also found that introns control intrinsic noise in both diploid and polyploid tissues. We found introns control intrinsic noise using a 5’-position dependent mechanism. Taken together, these results suggest that introns increase the probability of balanced, unbiased expression of autosomal alleles. This additional function of introns might explain why some genes have lost introns (e.g., diversity may be good for immune genes) and why most genes have retained introns. Intron control of intrinsic noise may also explain why some deep intronic mutations can result in allele silencing and are associated with human disease. Below, we show our results and then discuss the implications for cis control of intrinsic noise in metazoans.

## RESULTS

Here, we set out to study intrinsic noise *in vivo*, at the protein level, in the somatic tissues of the metazoan *Caenorhabditis elegans.* Specifically, we used reporter alleles controlled by the *hsp-90* promoter^26^ to conduct a survey of allele bias in *C. elegans* tissues. Figure 1 shows a diagram of differently colored reporter alleles encoding mCherry or mEGFP, and what high and low intrinsic noise would look like in the intestine tissue. A color-blind friendly version of the figure is available as Fig. S1. If alleles were expressed in a balanced biallelic fashion, then the color of the cells would be yellow, a merge of mEGFP (green) and mCherry (red). If there was a significant amount of allele bias, then cells would appear as color-shifted towards the color of the allele for which they are biased, shown in Figs. 1B&S1B. High noise animals would appear as a random “patch-work” of allelically biased and balanced cells.

**Fig. 1.**
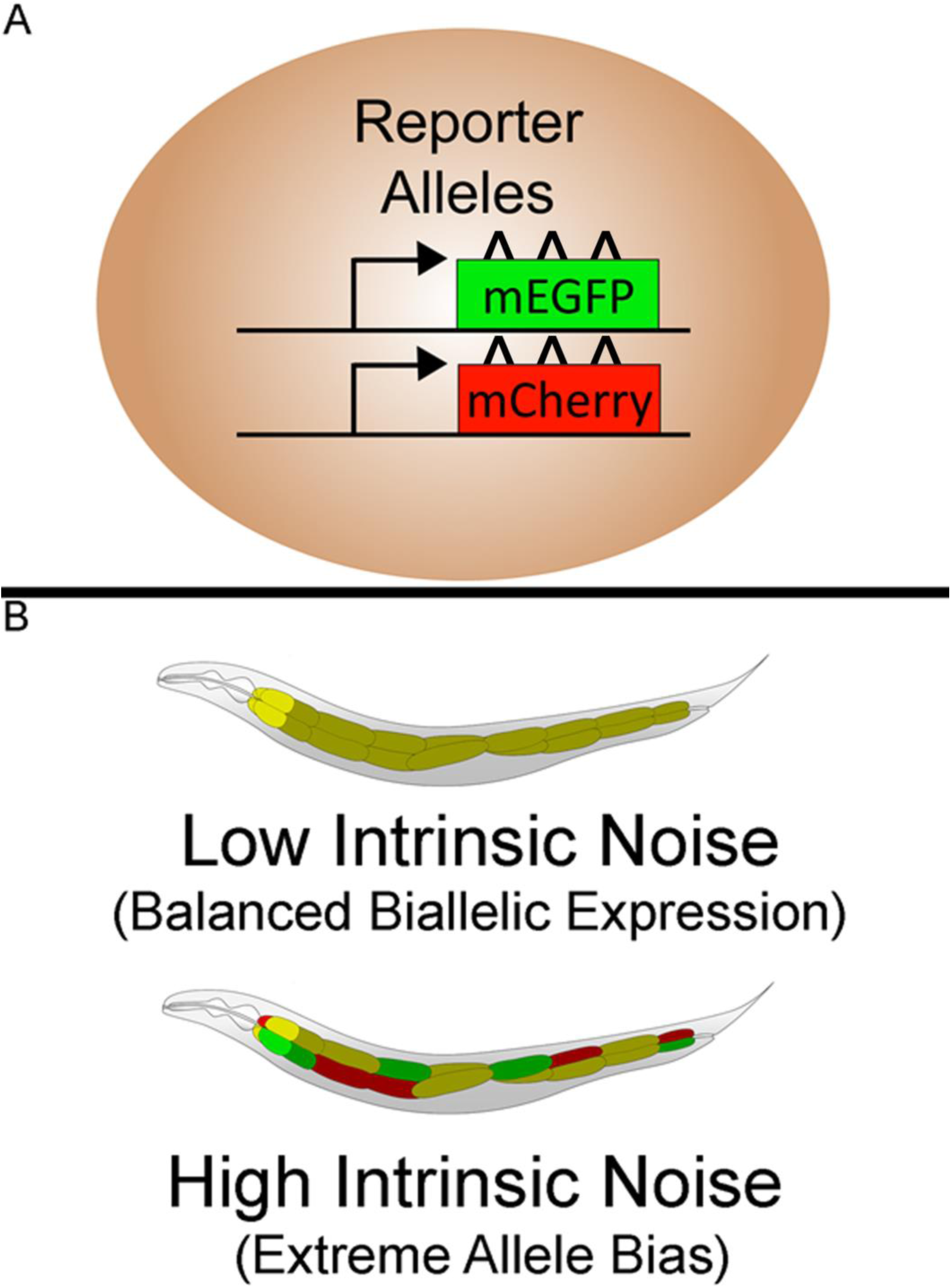
Experimental Schematics. **A)** shows a cartoon detailing that cells contain two otherwise identical copies of alleles encoding two spectrally distinct fluorescent proteins. **B)** shows cartoon examples of what high and low intrinsic noise would look like among the twenty cells comprising the worm intestine tissue. The top panel shows low intrinsic noise, which appear as yellow cells, because yellow is the balanced blend of red and green. The lower panel shows high intrinsic noise, which appears as an array of randomly biased and balanced cells; some cells appear yellow, and some cells are mostly, or all, red or green. Fig. S1 shows a colorblind friendly version of the cartoon in which green is switched to blue.

When we surveyed the tissues of day 2 adult *C. elegans*, scanning the heads and torsos of 40 individual worms, we observed striking allele bias in cells in several tissues, shown in Fig. 2. Note that these tissues are packed together in their native *interior milieu*^27^. Figure 2A shows a schematic cartoon overview of the worm. Figures 2B-H show representative microscopic images of various tissues surveyed. Figure 2B shows biased striated body wall muscles with the majority of other cells appearing yellow (biallelic). In the image, one can clearly observe that some muscle cells are skewed towards exclusively expressing the green or red allele. Figures 2C-E show higher magnification images of 4 neighboring muscle cells (labeled 1-4). Figure 2C shows the merged image, followed by the green channel (Fig. 2D) and red channel (Fig. 2E). For muscle cell number 2, notice that it appears red in the merged image and that the green signal is completely absent (mEGFP must have been absent from the cell or at a quantity below detection). In Figs. 2C-E, muscle cells 1&3 also appear to be monoallelic for the green allele, while muscle cell 4 is biallelic. Figures 2F-H show additional cells with extreme allele bias, including the excretory cell canal, a dorsal nerve cord, intestine cells, arcade cells and the smooth muscles in the pharynx.

**Fig. 2.**
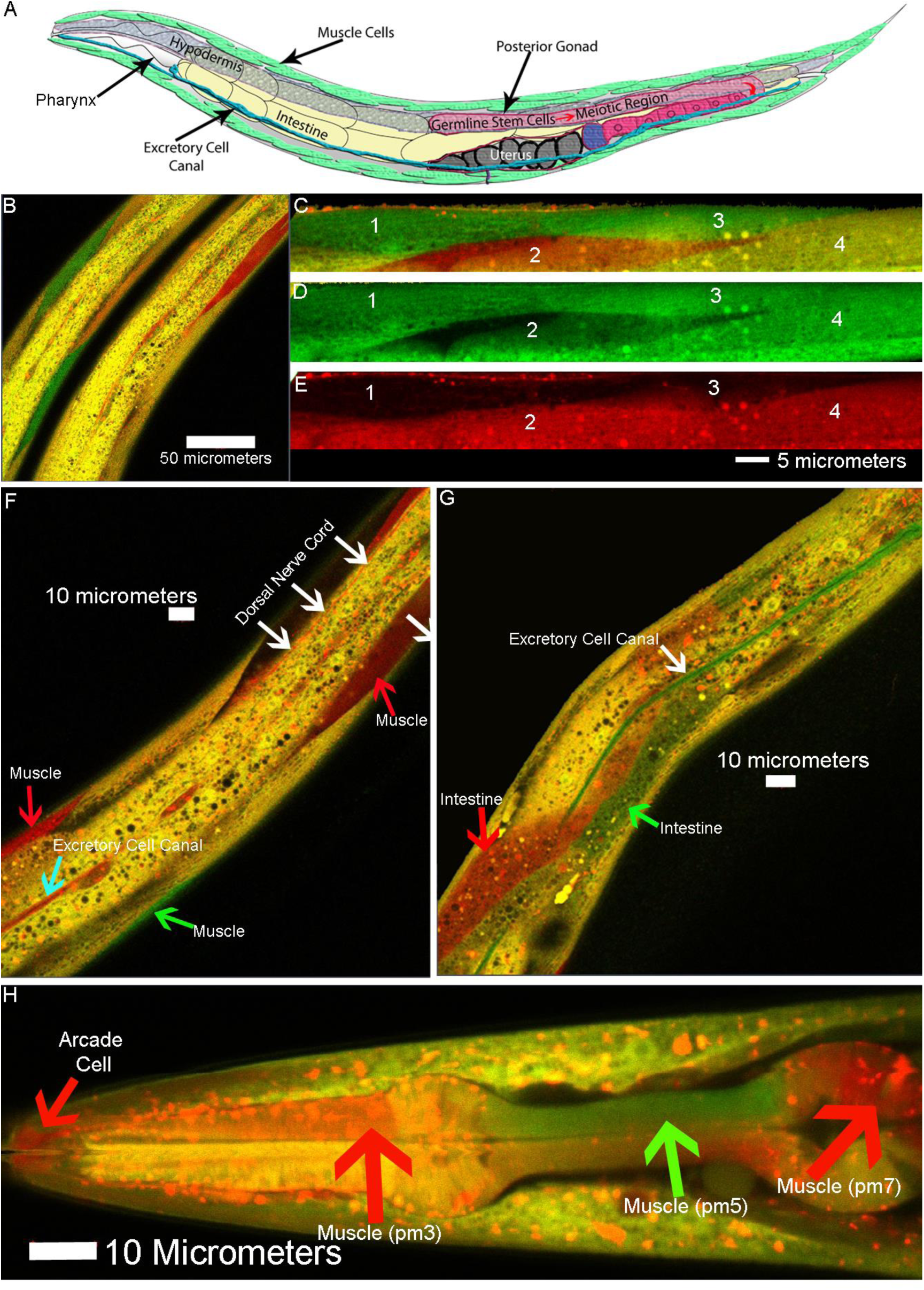
Allele bias can be fairly prevalent in somatic tissues of adult *C. elegans*. **A)** shows a schematic overview of *C. elegans* anatomy. **B)** shows merged, two-color confocal micrographs of “torso” of two animals expressing differently colored reporter alleles, showing monoallelic expression/allele bias in muscle cells; anterior/head is up, hypodermis is the yellow strip in the center of each animal. **C)** shows a composite confocal micrograph of four body wall muscle cells from an individual animal, each numbered 1-4. **D&E** show the individual mEGFP and mCherry channels. Cells 1, 2 & 3 were monoallelic (or really extreme), and cell 4 expressed both alleles. **F)** shows another animal with allele bias/monoallelic expression in muscle cells, a dorsal nerve cord and the excretory cell canal. **G)** shows another animal with allele bias in muscle cells, intestine cells and the excretory cell canal. **H)** shows the head of an animal with monoallelic expression/allele bias in its smooth pharynx muscles and the structural arcade cells. Red puncta are lysosomes, delineated by concentrated, acid/protease resistant, mCherry ^4,50^. Images with clear bias and good cell positioning were selected from four independent experiments capturing z stack image sets of ten two day old adult animals in each experiment.

The *hsp-90* reporter alleles used for the tissue survey contained fluorescent reporter genes with the same three synthetic introns typically used in *C. elegans* reporters^28-30^. We recently showed that these introns increase gene expression levels in *C. elegans* using a conserved 5’-proximal intron position dependent mechanism^31^. In 2012, another report detailed that 5’-introns recruit chromatin marks associated with (or causative of) an active chromatin state^25^. The idea that emerged from considering these two studies is that, if introns open chromatin and increase expression levels, then introns should also increase the probability that both alleles of a gene are expressed. Therefore, we tested the hypothesis that introns affect intrinsic noise. To test our hypothesis, we engineered worms to express reporter alleles that were identical to the *hsp-90* alleles used in the above survey, but we removed introns from the coding sequence of the fluorescent proteins (Fig 3A). We then measured the expression levels of the intron-bearing and intronless sets of reporter alleles, which were expressed from the exact same locus (tt5605 on Chromosome II), using a confocal microscope (Fig. 3B), as we described previously^32^. We quantified allele bias in individual worm intestine cells as intrinsic noise, visible as deviation from the 45° central trendline on scatter plots, using the same analytical framework as prior reports quantifying intrinsic noise in bacteria^9^ and yeast^8^ cells.

**Fig. 3.**
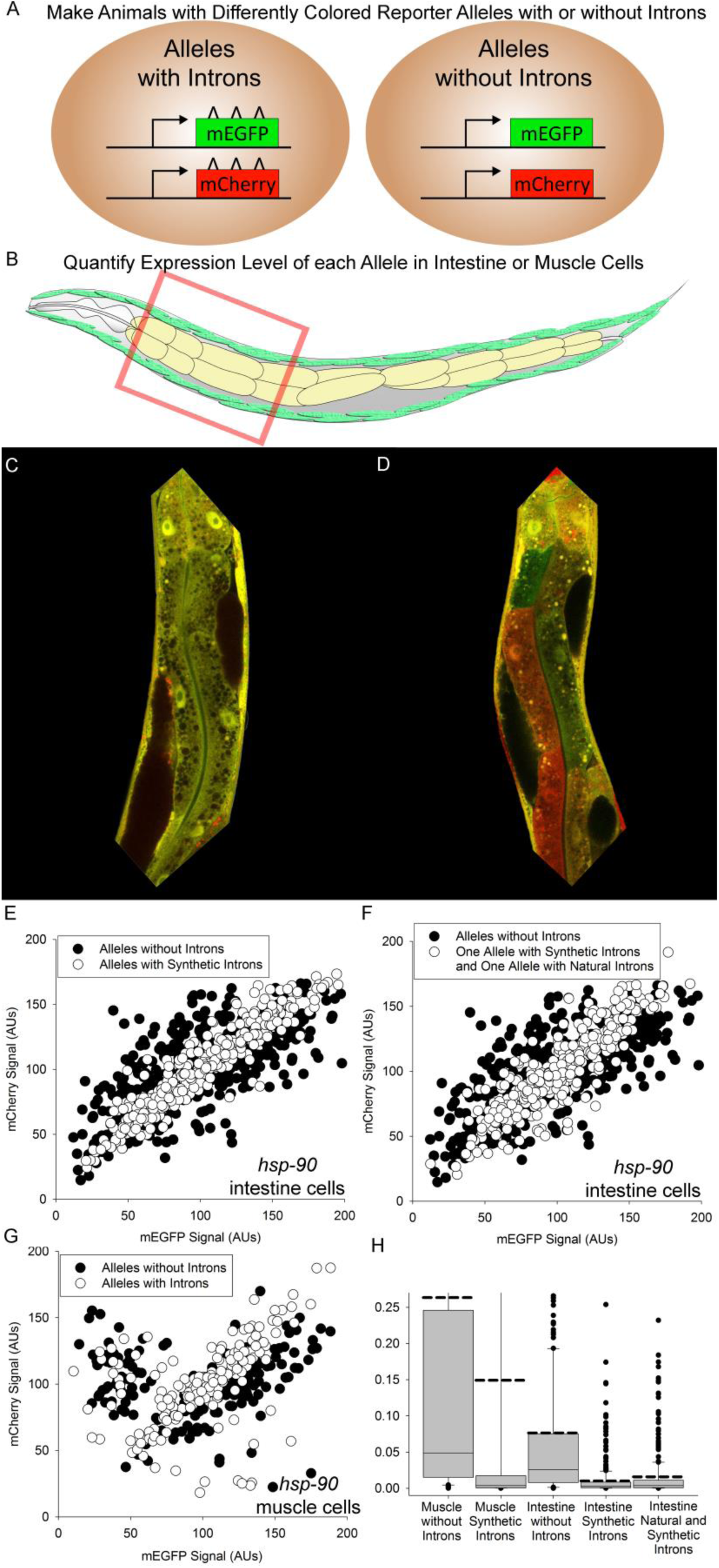
Schematic overview of experimental design and effects of introns on intrinsic noise. All plots use arbitrary units (AUs). **A)** shows a schematic of reporter allele design. **B)** shows the field of view we used to measure allele expression levels in muscle and intestine cells. **C)** shows an image of an animal’s intestine in which alleles have introns. **D)** shows an image of an animal’s intestine in which alleles do not have introns. **E)** shows a scatter plot of allele expression levels in intestine cells; black dots also underlie white dots. Raw data in Data S1. **F)** shows a scatter plot of allele expression levels in intestine cells; black dots also underlie white dots. Raw data in Data S1. **G)** shows a scatter plot of allele expression levels in muscle cells; black dots also underlie white dots. Raw data in Data S1. **H)** shows box plots of intrinsic noise; solid line is median, dashed line is mean, dots are outliers. A full scale version of Fig. 2H showing outliers is available as Fig. S2. Hundreds of cells were sampled across at least three independent experiments measuring cells from at least ten individual animals per group for each experiment type. Additional statistical details available in Supplementary Information.

Figure 3C shows a representative image of the anterior half of an intestine, from an animal that expresses intron-bearing alleles. Notice that intestine cells appear yellow, due to biallelic expression. Figure 3D shows a representative image of another anterior intestine, from an animal that expresses intronless reporter alleles. In this animal, some cells appear yellow due to biallelic expression, while others are skewed towards either the red or green allele. Additional images are found in Fig. 4B. Figure 3E shows the scatter plots from this experiment, with intron containing alleles represented by white dots and intronless alleles represented by black dots. More deviation from the 45° trendline indicates more intrinsic noise. Some cells expressing intron-bearing alleles (white dots) deviate relatively far from the 45° trendline, but most do not, indicating relatively low intrinsic noise. However, the intronless alleles (black dots) deviate further and more frequently, surrounding the intron-bearing alleles in the scatter plot. In fact, the removal of synthetic introns from alleles significantly increased median intrinsic noise in gene expression by approximately an order of magnitude, from 0.00275 to 0.0255, shown in Fig. 3H (*P* < 0.05; See Supplementary Information for additional details like Q scores). We found the same effect when one of the reporter alleles contained two natural introns found in *hsp-90*, shown in Fig. 3H (median intrinsic noise from 0.00376 to 0.0255; *P* < 0.05). Thus, the presence of introns in the coding sequence of these alleles reduced stochastic allele biases, thereby increasing the probability of a biallelic state of gene expression. This previously undescribed function for introns may be important for ensuring the high level of coordinated gene expression that multicellular life requires. We call this <u>i</u>ntron-<u>co</u>ntrol of intrinsic <u>n</u>oise (ICON).

**Fig. 4.**
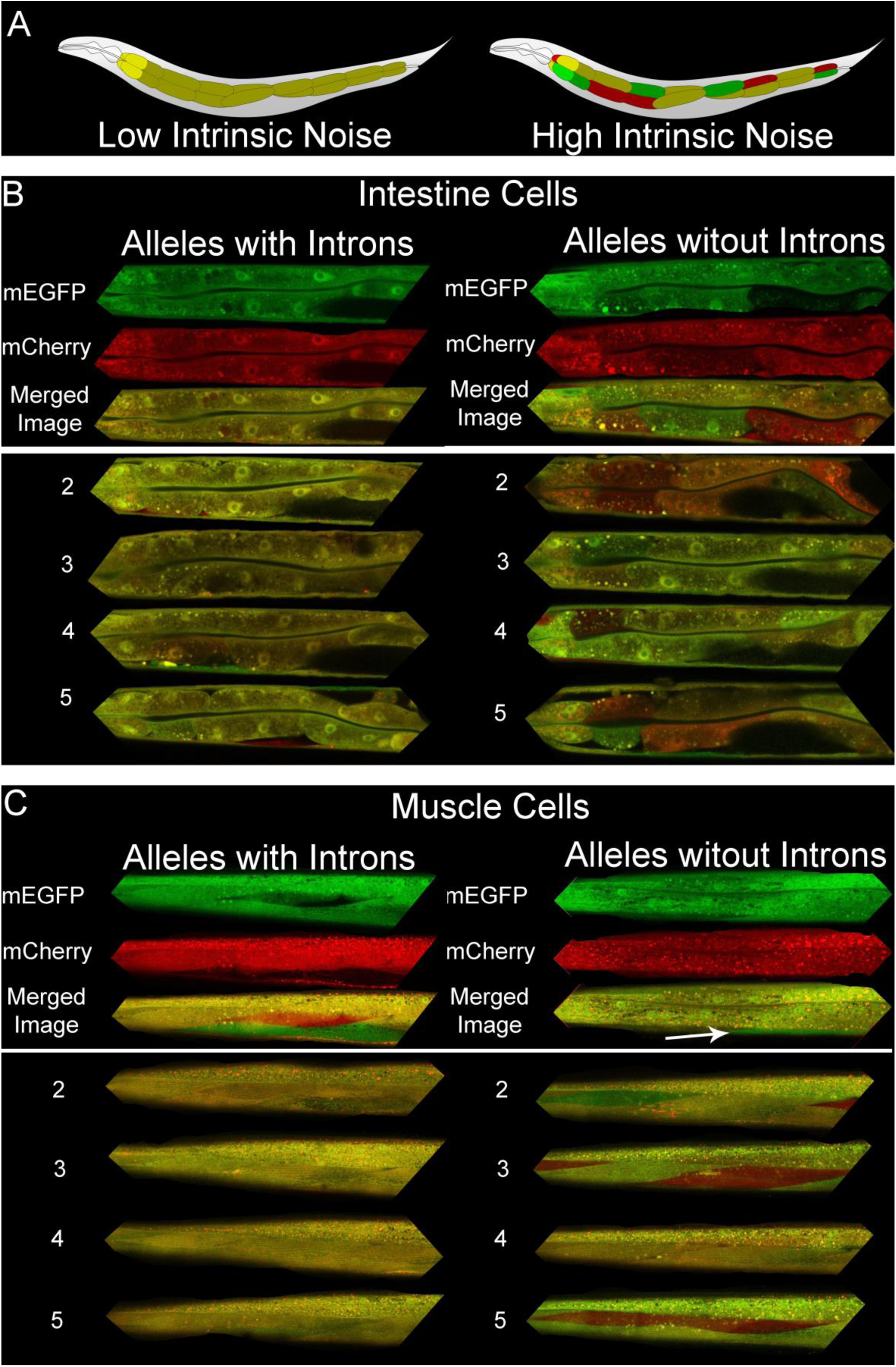
Cartoon schematics and microscopic images of alleles with and without introns in intestine and muscle tissues. **A)** shows cartoon images of what high and low intrinsic noise in intestine cells would look like. Images in **B&C** are fluorescent micrographs showing expression of *hsp-90* controlled reporter alleles encoding mCherry or mEGFP in individual cells in intact intestine and muscle tissues. Anterior is left, and dorsal is top. The top of each image contains a dotty swath of adjacent hypodermal tissue. The first representative animal in each group shown is displayed in individual and merged microscopic channels, and the tissues of the subsequent individual animals, labeled 2-5 are shown as merged microscopic images. **B)** shows expression of reporter alleles with and without introns in polyploid intestine cells. **C)** shows expression of reporter alleles with and without introns in diploid striated muscle cells. We arbitrarily selected images from the images underlying the data Fig.3 in which animals and their tissues were better oriented for viewing the cell types with minimal distortion by other tissues or oblique viewing angles. This tends to be more difficult for the muscle cells because *C. elegans* prefer to orient their lateral faces towards the ground, not unlike a flounder. And, in the top right series of images of muscles from animals expressing intronless alleles, **t**he hypodermal swath of issue is largely in view, and a white arrow points to a virtually monoallelic green cell, oriented obliquely.

Genes are often regulated in a tissue specific manner, and chromatin signature based evidence suggests that this is true for monoallelic expression as well^20,33,34^. Specifically, at the monoallelic gene expression database, some genes appear to have a monoallelic chromatin signature in some but not all tissues investigated. These reports suggest that intrinsic noise for a given gene may also be tissue specific in *C. elegans*. Additionally, intrinsic noise should be higher in diploid tissues, relative to polyploid tissues, because polyploid tissues have more chances (copies of the diploid genome) to express both alleles. To answer these questions, we used the same *hsp-90* reporter alleles described above to study intrinsic noise in diploid muscle cells.

Similar to what we see in intestine cells, muscle cells that express intron-bearing alleles appear mostly yellow, indicating biallelic expression, though some cells are extremely biased (Fig. 4C). Animals expressing intronless alleles showed a far more variegated gene expression pattern (Fig. 4C). When we quantified intrinsic noise in both sets of animals, we found that introns decreased median intrinsic noise in muscles by more than an order of magnitude, from a median value of 0.0489 to 0.00365 (Fig. 3G; *P* < 0.05).

When we combined data from all of our experiments in intestine and muscle cells, we found that intrinsic noise was significantly higher in (2n) muscle cells than in (16n) polyploid intestines (Fig. 3H; *P* < 0.001; One Way ANOVA with Dunn’s for all alleles together). This result is expected, given the increased chance for biallelic expression in polyploid cells. However, when we looked at just intron-bearing alleles, the difference was not significant (*P =* 0.11 in One-Way ANOVA with Dunn’s test; Power below 0.8; not a confident negative result). Thus, the tissue-specific difference in intrinsic noise for all *hsp-90* alleles together is driven by the intronless alleles. This is likely due to the differences in intrinsic noise set points between intronless and intron-bearing allele pairs. Since intron-bearing alleles are mainly biallelic (low intrinsic noise set point), there is a relatively small decrease in noise that is possible, indicating the strong influence introns have in restricting seemingly stochastic, cell autonomous allele biases.

Intrinsic noise is known to be controlled by promoter sequence in *Saccharomyces cerevisiae*^8^. To determine if ICON was affected by promoter in metazoans, we measured the same reporter alleles with and without introns, controlled by two additional promoters. We tested the promoter from *vit-2*, which encodes intestinally expressed yolk protein, and is perhaps the strongest promoter in *C. elegans*^35^. For these allele pairs, we found that introns significantly decreased median intrinsic noise by about 82%, from 0.00722 to 0.00133 (Figs. 5A&C; *P* < 0.05). We also tested a distinctly regulated, inducible, small heat shock protein promoter from *hsp-16.2*^36,37^. In animals expressing *hsp-16.2* reporter alleles, the presence of introns decreased intrinsic noise by about 50%, from 0.00931 to 0.00417 (Figs. 5B&C; *P* < 0.05). While introns restricted intrinsic noise under the control of distinct promoters, the effect sizes were different, demonstrating that promoters also affected allele bias. Our finding that each gene can have a gene-specific quantity of (or set-point for) intrinsic noise is in agreement with different promoters having different amounts of intrinsic noise in diploid yeast^8^. Statistical comparisons of alleles controlled by different promoters with and without introns or all together are shown in Supplements (Statistical Analyses, pages 5&6).

**Fig. 5.**
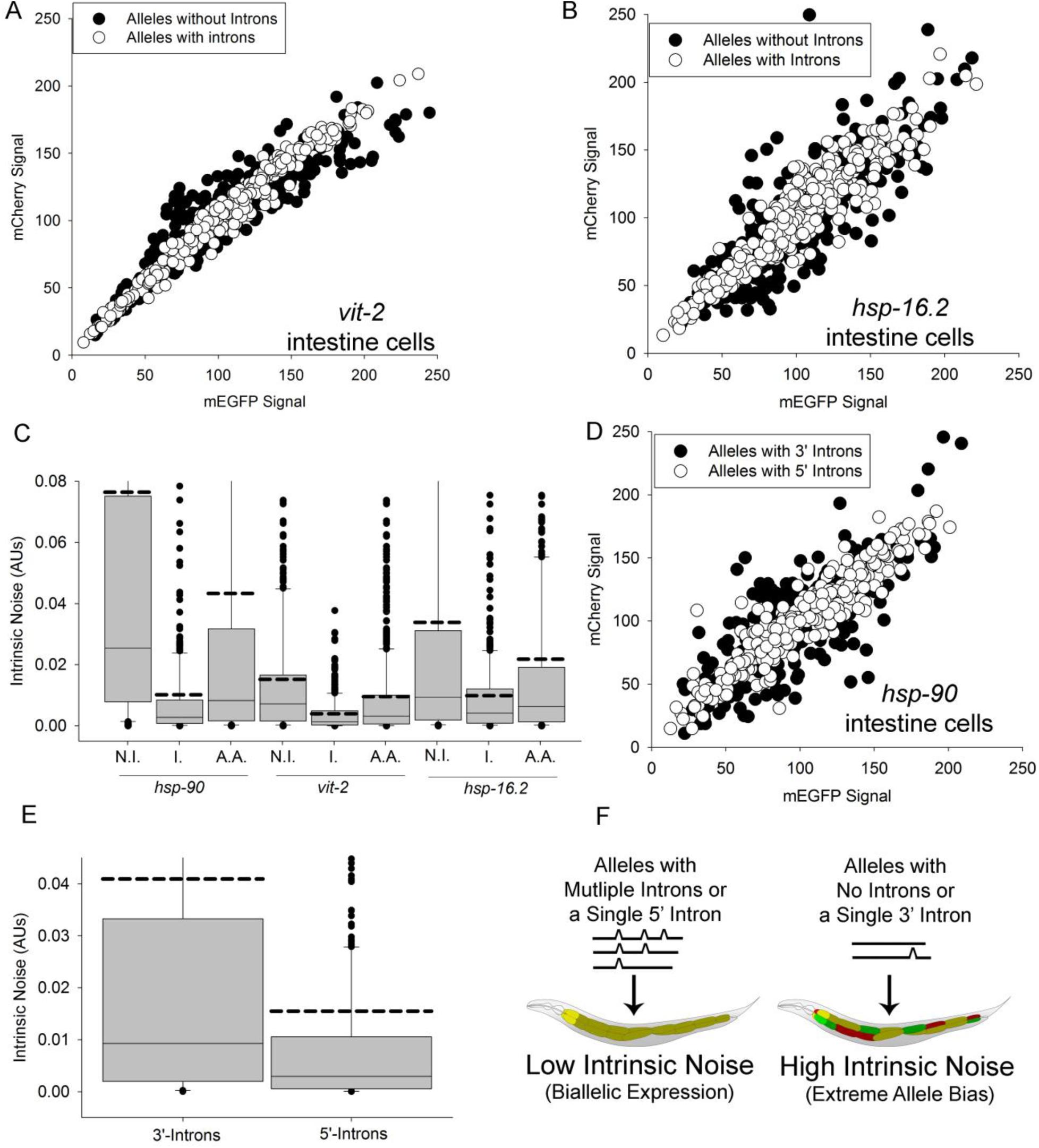
Introns control intrinsic noise with distinctly regulated promoters and use a position dependent mechanism for ICON. All plots use arbitrary units (AUs). **A)** shows a scatter plot of *vit-2* reporter alleles in intestine cells; black dots also underlie white dots. Raw data in Data S1. **B)** shows a scatter plot of *hsp-16.2* reporter alleles in intestine cells; black dots also underlie white dots. Raw data in Data S1. **(C)** shows boxplots of intrinsic noise levels for each different promoter, for alleles with no introns (N.I.), for alleles with introns (I.), and for all alleles combined (A.A.); solid line is median, dashed line is mean. Extended graph showing full range and outliers shown in Fig. S3. **D)** shows a scatterplot of reporter alleles with single introns; black dots also underlie white dots. Raw data in Data S1. **E)** shows the boxplots of intrinsic noise for reporter alleles; solid line is median, dashed line is mean. Extended graph showing full range and outliers shown in Fig. S4. Hundreds of cells were sampled across at least four independent experiments measuring cells from at least ten individual animals per group for each experiment type. Additional statistical details available in Supplementary Information. **F)** shows a cartoon diagram illustrating the major findings of this study.

Next, because introns close to the TSS are associated with open chromatin markings^25^, we hypothesized that the same 5’-position dependent mechanism used for IME^31^ may also be used for ICON. To test this, we created reporter allele pairs with a single intron positioned close to the transcription start site, or positioned more distally toward the 3’UTR (here, called 5’-intron or 3’-intron, respectively). Worms that expressed alleles with 5’-introns had balanced biallelic expression, similar to the worms that expressed alleles with multiple introns. Worms expressing alleles with single 3’-introns showed a variegated allele expression patterns, similar to intronless worms. Alleles with single 5’-introns had significantly less intrinsic noise than alleles with single 3’-introns (0.00296 vs .00928 for median intrinsic noise; *P* < 0.05), shown in Fig. 5E. Thus, the mechanism by which introns control intrinsic noise is intron-position dependent. Fig. 5F shows a graphical illustration of the major findings of this study.

About 3% of *C. elegans’* protein-coding genes are intronless (Data S2). Of the 20,390 protein-coding genes in the human genome (GRCh38), 1,165 are intronless – about 6% (Data S2). We performed GO terms enrichment analyses on human intronless genes and found many immune activity-related GO terms (Data S2). We also found enrichment for “G protein coupled receptor activity” and “olfactory receptor activity”. Olfactory receptors are known to be expressed in an exclusively monoallelic fashion^38^. Over 500 of the intronless genes are listed as monoallelic in the database for monoallelic expression (dbMAE^33^; Data S2). It is possible some genes lost introns to ensure variegated allele expression. It now seems plausible that many metazoan genes retain introns because they decrease intrinsic noise. Put differently, introns increase the chance for equal expression of both parental alleles in a cell. Intron control of intrinsic noise may be one additional reason why many metazoan genes have introns, especially those genes that retain single, relatively 5’ introns and have no alternative splicing isoforms.

## DISCUSSION

Previous reports analyzed intrinsic noise in bacteria^9^, and in diploid yeast^8^. For these organisms, intrinsic noise is higher by about an order of magnitude or more than what we have measured here in the metazoan, *C. elegans*. Intrinsic noise appears to be more restricted in metazoans. Most genes in the human genome, and in most metazoan genomes for that matter, have introns. Considering our results, it seems likely that one reason to have introns is to ensure coordinated and robust gene expression in multicellular organisms. In metazoans, like humans, the question remains as to why some genes have lost their introns^39^. For some classes of genes, like chaperones, the loss seems attributable to poor splicing at higher temperatures. For other classes of genes, loss of introns may occur to ensure variegated allele expression. For example, G-protein coupled receptors (GPCRs) have lost introns at a relatively increased rate ^39^. Variegation of cell surface receptor alleles may be advantageous for ensuring some cells remain uninfected by a virus (Red Queen hypothesis reviewed in^40^).

Single, 5’-positioned introns are sufficient for IME^41^ presumably because, among other reasons, they recruit chromatin opening marks^25^. Our data suggest that this same mechanism is shared with ICON. However, the data for effect sizes of ICON compared to IME suggest that there may also be mechanistic components that are not shared between these phenomena. We previously showed that IME effect size was similar under control of the *vit-2* and *hsp-90* promoters (about 50%)^31^. Here, using the same promoters, we found that *vit-2* and *hsp-90* had different ICON effect sizes (50% vs. an approximate order of magnitude in the intestine cells). Thus, while promoter had little influence on IME, it had a large influence on ICON. This difference suggests that the underlying mechanisms controlling IME and ICON may not be entirely the same.

Our experimental system showed that intrinsic noise can be a significant component of cell-to-cell variation in gene expression *in vivo* in the metazoan *C. elegans.* These results are significant because they demonstrate that intrinsic noise can be observed in an experimentally tractable metazoan *in vivo*, corroborating previous *in vitro* studies^17,21,42,43^. Additionally, these results are significant because they allow us to examine intrinsic noise at the protein level, complementing RNA based studies. In some scenarios, there can be little correlation between protein and RNA levels, as in^44^, and reviewed in 2012 here^45^ and in 2016 here^46^. Additionally, studies of allele bias at the RNA level can sometimes have trouble distinguishing between transcriptional bursting^47^ and actual allele bias that manifests at the protein level^13^. Detecting allele specific expression at the protein level by means other than those described here remain challenging. For example, it can be technically difficult or impossible to generate antibodies that distinguish between two allelic variants of the same gene. We anticipate that this experimental system can be used to identify trans acting factors that control intrinsic noise.

RNA and chromatin marking based evidence suggests that extreme allele bias or monoallelic gene expression is prevalent in murine and human cells^16-21^. Furthermore, monoallelic expression of particular genes could explain some perplexing clinical scenarios. For example, some people harbor dominant negative oncogenes, but remain cancer free^1,2^. It would be interesting to monitor healthy tissue in these scenarios to determine if the dominant negative allele is being silenced. In another scenario, fortunate monoallelic expression of a “good” allele protected some, but not all, family members from the negative consequences of a dominant negative *PIT1* allele^48^. In this study, the father and grandmother of the affected patient harbored the dominant negative allele, but no mRNA of that allele was detected, suggesting that they were protected by allelic silencing. Finally, an intriguing clinical case of a single patient with COL6A2-associated Bethlem myopathy suggests that intronic mutations can lead to silencing and potential disease^49^. In this study, the affected patient harbored one allele with a large intronic mutation and a second allele with a mutation in exon 28 that is predicted to cause disease. The allele with the intronic mutation was silenced, leading to the sole expression of the potential disease allele. Taken together, these cases demonstrate that extreme allele bias can be consequential, and support the idea that introns increase the probability of a balanced, biallelic state of gene expression. More work is necessary to determine how different lengths and sequences of introns, or sequences within introns, including clinically relevant mutations, influence intrinsic noise.

## Acknowledgments

We would like to thank Gary Ruvkun, Chris Link, George Martin and Matt Kaeberlein for careful reading of manuscript drafts. We would like to thank Lu Wang, Theo Bammler and James MacDonald at the University of Washington Nathan Shock Center for Excellence in the Basic Biology of Aging for identifying intronless genes and GO Terms analyses.

## Author contributions

AM and BS designed the study. BS performed microscopy and image cytometry. SY performed crosses and animal husbandry. BS, AM and SY performed molecular cloning, microinjections and animal husbandry to make transgenic animals. BS and AM analyzed the data. BS and AM wrote the initial manuscript. BS, SY and AM revised the manuscript.

## Data and materials availability

All data is available in the main text or the Supplementary Information.

## Supplementary Materials

Supplementary Information, containing: Detailed Statistical Testing Results, Figures S1-S4 & Tables S1-S2 & Materials and Methods.

Data S1, containing raw data.

Data S2, containing results of bioinformatic analyses.

