## Supplementary Information for "Introns Control Stochasticity in Metazoan Gene Expression"

**This PDF file includes:**

Statistical Analyses……………………………………………………………Pages 2-7
Figs. S1 to S4…………………………………………………………...……Pages 8-11
Tables S1 to S2…………………………………………………………...Pages 12-13
Caption for Data S1…..…………………………………………………………Page 14
Caption for Data S2…..…………………………………………………………Page 14
Materials and Methods…………………………………………………………Pages 15-18

**Other Supplemental Materials for this manuscript include the following:**

Data S1
Data S2

### Supplemental Text

_Results of Statistical Analyses_

Below, we detail the results of each statistical analysis of our data on intrinsic noise levels.

**Kruskal-Wallis One Way Analysis of Variance on Ranks for hsp-90 reporter alleles**

| Group | N | Missing | Median | 25% | 75% |
|---|---|---|---|---|---|
| Muscle No Introns Noise | 180 | 0 | 0.0489 | 0.0151 | 0.246 |
| Muscle Introns Noise | 180 | 0 | 0.00365 | 0.000603 | 0.0174 |
| Gut Cell No Introns Noise | 398 | 0 | 0.0255 | 0.00786 | 0.0751 |
| Gut Cell Introns Noise | 398 | 0 | 0.00275 | 0.000802 | 0.00846 |
| Gut Cell Hybrid Noise | 380 | 0 | 0.00376 | 0.000694 | 0.0111 |

H = 376.768 with 4 degrees of freedom. (P = <0.001)

The differences in the median values among the treatment groups are greater than would be expected by chance; there is a statistically significant difference (P = <0.001)

To isolate the group or groups that differ from the others use a multiple comparison procedure.

All Pairwise Multiple Comparison Procedures (Dunn's Method) :

| Comparison | Diff of Ranks | Q | P<0.05 |
|---|---|---|---|
| Muscle No Int vs Gut Cell Intr | 577.110 | 14.485 | Yes |
| Muscle No Int vs Gut Cell Hybr | 539.015 | 13.431 | Yes |
| Muscle No Int vs Muscle Intron | 476.594 | 10.194 | Yes |
| Muscle No Int vs Gut Cell No I | 148.155 | 3.719 | Yes |
| Gut Cell No I vs Gut Cell Intr | 428.955 | 13.643 | Yes |
| Gut Cell No I vs Gut Cell Hybr | 390.859 | 12.286 | Yes |
| Gut Cell No I vs Muscle Intron | 328.439 | 8.244 | Yes |
| Muscle Intron vs Gut Cell Intr | 100.516 | 2.523 | No |
| Muscle Intron vs Gut Cell Hybr | 62.420 | 1.555 | Do Not Test |
| Gut Cell Hybr vs Gut Cell Intr | 38.096 | 1.198 | Do Not Test |

Note: The multiple comparisons on ranks do not include an adjustment for ties.

**Kruskal-Wallis One Way Analysis of Variance on Ranks for Promoters and Introns**

| Group | N | Missing | Median | 25% | 75% |
|---|---|---|---|---|---|
| vit2 introns noise | 359 | 0 | 0.00133 | 0.000281 | 0.00491 |
| hsp90 introns noise | 398 | 0 | 0.00275 | 0.000802 | 0.00846 |
| hsp162 introns noise | 360 | 0 | 0.00417 | 0.000869 | 0.0121 |
| vit2 no introns noise | 351 | 0 | 0.00722 | 0.00160 | 0.0166 |
| hsp90 no introns noise | 398 | 0 | 0.0255 | 0.00786 | 0.0751 |
| hsp162 no introns noise | 357 | 0 | 0.00931 | 0.00190 | 0.0312 |

H = 430.516 with 5 degrees of freedom. (P = <0.001)

The differences in the median values among the treatment groups are greater than would be expected by chance; there is a statistically significant difference (P = <0.001)

To isolate the group or groups that differ from the others use a multiple comparison procedure.

All Pairwise Multiple Comparison Procedures (Dunn's Method) :

| Comparison | Diff of Ranks | Q | P<0.05 |
|---|---|---|---|
| hsp90 no intr vs vit2 introns | 876.368 | 18.758 | Yes |
| hsp90 no intr vs hsp90 introns | 663.075 | 14.573 | Yes |
| hsp90 no intr vs hsp162 intron | 580.284 | 12.429 | Yes |
| hsp90 no intr vs vit2 no intro | 438.962 | 9.340 | Yes |
| hsp90 no intr vs hsp162 no int | 297.525 | 6.359 | Yes |
| hsp162 no int vs vit2 introns | 578.843 | 12.065 | Yes |
| hsp162 no int vs hsp90 introns | 365.550 | 7.813 | Yes |
| hsp162 no int vs hsp162 intron | 282.759 | 5.898 | Yes |
| hsp162 no int vs vit2 no intro | 141.437 | 2.931 | No |
| vit2 no intro vs vit2 introns | 437.406 | 9.078 | Yes |
| vit2 no intro vs hsp90 introns | 224.114 | 4.768 | Yes |
| vit2 no intro vs hsp162 intron | 141.323 | 2.935 | No |
| hsp162 intron vs vit2 introns | 296.084 | 6.184 | Yes |
| hsp162 intron vs hsp90 introns | 82.791 | 1.773 | No |
| hsp90 introns vs vit2 introns | 213.292 | 4.565 | Yes |

Note: The multiple comparisons on ranks do not include an adjustment for ties.

**Kruskal-Wallis One Way Analysis of Variance on Ranks for Promoters with All Alleles**

| Group | N | Missing | Median | 25% | 75% |
|---|---|---|---|---|---|
| vit-2 | 710 | 0 | 0.00321 | 0.000625 | 0.0100 |
| hsp90 | 796 | 0 | 0.00828 | 0.00165 | 0.0317 |
| hsp16 | 717 | 0 | 0.00631 | 0.00130 | 0.0191 |

H = 100.947 with 2 degrees of freedom. (P = <0.001)

The differences in the median values among the treatment groups are greater than would be expected by chance; there is a statistically significant difference (P = <0.001)

To isolate the group or groups that differ from the others use a multiple comparison procedure.

All Pairwise Multiple Comparison Procedures (Dunn's Method) :

| Comparison | Diff of Ranks | Q | P<0.05 |
|---|---|---|---|
| hsp90 vs vit-2 | 328.591 | 9.917 | Yes |
| hsp90 vs hsp16 | 107.959 | 3.267 | Yes |
| hsp16 vs vit-2 | 220.633 | 6.492 | Yes |

Note: The multiple comparisons on ranks do not include an adjustment for ties.

**Kruskal-Wallis One Way Analysis of Variance on Ranks for All Possible Promoter Comparisons**

| Group | N | Missing | Median | 25% | 75% |
|---|---|---|---|---|---|
| vit2 introns noise | 359 | 0 | 0.00133 | 0.000281 | 0.00491 |
| hsp90 Introns Noise | 398 | 0 | 0.00275 | 0.000802 | 0.00846 |
| hsp162 introns noise | 360 | 0 | 0.00417 | 0.000869 | 0.0121 |
| vit2 no introns noise | 351 | 0 | 0.00722 | 0.00160 | 0.0166 |
| hsp90 no introns noise | 398 | 0 | 0.0255 | 0.00786 | 0.0751 |
| hsp162 no introns noise | 357 | 0 | 0.00931 | 0.00190 | 0.0312 |
| vit-2 | 710 | 0 | 0.00321 | 0.000625 | 0.0100 |
| hsp90 | 796 | 0 | 0.00828 | 0.00165 | 0.0317 |
| hsp16 | 717 | 0 | 0.00631 | 0.00130 | 0.0191 |

H = 531.584 with 8 degrees of freedom. (P = <0.001)

The differences in the median values among the treatment groups are greater than would be expected by chance; there is a statistically significant difference (P = <0.001)

To isolate the group or groups that differ from the others use a multiple comparison procedure.

All Pairwise Multiple Comparison Procedures (Dunn's Method) :

| Comparison | Diff of Ranks | Q | P<0.05 |
|---|---|---|---|
| hsp90 no intr vs vit2 introns | 1752.736 | 18.760 | Yes |
| hsp90 no intr vs hsp90 Introns | 1326.151 | 14.574 | Yes |
| hsp90 no intr vs vit-2 | 1320.258 | 16.426 | Yes |
| hsp90 no intr vs hsp162 intron | 1160.569 | 12.431 | Yes |
| hsp90 no intr vs hsp16 | 878.992 | 10.955 | Yes |
| hsp90 no intr vs vit2 no intro | 877.924 | 9.341 | Yes |
| hsp90 no intr vs hsp90 | 663.075 | 8.415 | Yes |
| hsp90 no intr vs hsp162 no int | 595.050 | 6.360 | Yes |
| hsp162 no int vs vit2 introns | 1157.686 | 12.067 | Yes |
| hsp162 no int vs hsp90 Introns | 731.101 | 7.814 | Yes |
| hsp162 no int vs vit-2 | 725.208 | 8.708 | Yes |
| hsp162 no int vs hsp162 intron | 565.519 | 5.899 | Yes |
| hsp162 no int vs hsp16 | 283.942 | 3.415 | Yes |
| hsp162 no int vs vit2 no intro | 282.874 | 2.932 | No |
| hsp162 no int vs hsp90 | 68.025 | 0.832 | Do Not Test |
| hsp90 vs vit2 introns noise | 1089.660 | 13.353 | Yes |
| hsp90 vs hsp90 Introns Noise | 663.075 | 8.415 | Yes |
| hsp90 vs vit-2 | 657.183 | 9.918 | Yes |
| hsp90 vs hsp162 introns noise | 497.493 | 6.102 | Yes |
| hsp90 vs hsp16 | 215.917 | 3.267 | Yes |
| hsp90 vs vit2 no introns noise | 214.848 | 2.612 | Do Not Test |
| vit2 no intro vs vit2 introns | 874.812 | 9.079 | Yes |
| vit2 no intro vs hsp90 Introns | 448.227 | 4.769 | Yes |
| vit2 no introns noise vs vit-2 | 442.335 | 5.281 | Yes |
| vit2 no intro vs hsp162 intron | 282.645 | 2.936 | No |
| vit2 no introns noise vs hsp16 | 1.069 | 0.0128 | Do Not Test |
| hsp16 vs vit2 introns noise | 873.743 | 10.528 | Yes |
| hsp16 vs hsp90 Introns Noise | 447.158 | 5.573 | Yes |
| hsp16 vs vit-2 | 441.266 | 6.493 | Yes |
| hsp16 vs hsp162 introns noise | 281.576 | 3.396 | Do Not Test |
| hsp162 intron vs vit2 introns | 592.167 | 6.185 | Yes |
| hsp162 intron vs hsp90 Introns | 165.582 | 1.774 | No |

| | | | |
|---|---|---|---|
| hsp162 introns noise vs vit-2 | 159.690 | 1.923 | Do Not Test |
| vit-2 vs vit2 introns noise | 432.477 | 5.203 | Yes |
| vit-2 vs hsp90 Introns Noise | 5.893 | 0.0733 | Do Not Test |
| hsp90 Introns vs vit2 introns | 426.585 | 4.566 | Yes |

Note: The multiple comparisons on ranks do not include an adjustment for ties.

**Mann-Whitney Rank Sum Test for Alleles with single 5' or 3' introns**

**Normality Test (Shapiro-Wilk)** Failed (P < 0.050)

| Group | N | Missing | Median | 25% | 75% |
|---|---|---|---|---|---|
| 3' noise | 360 | 0 | 0.00928 | 0.00198 | 0.0333 |
| 5' noise | 360 | 0 | 0.00296 | 0.000562 | 0.0106 |

Mann-Whitney U Statistic= 45006.000

T = 149574.000 n(small)= 360 n(big)= 360 (P = <0.001)

The difference in the median values between the two groups is greater than would be expected by chance; there is a statistically significant difference (P = <0.001).

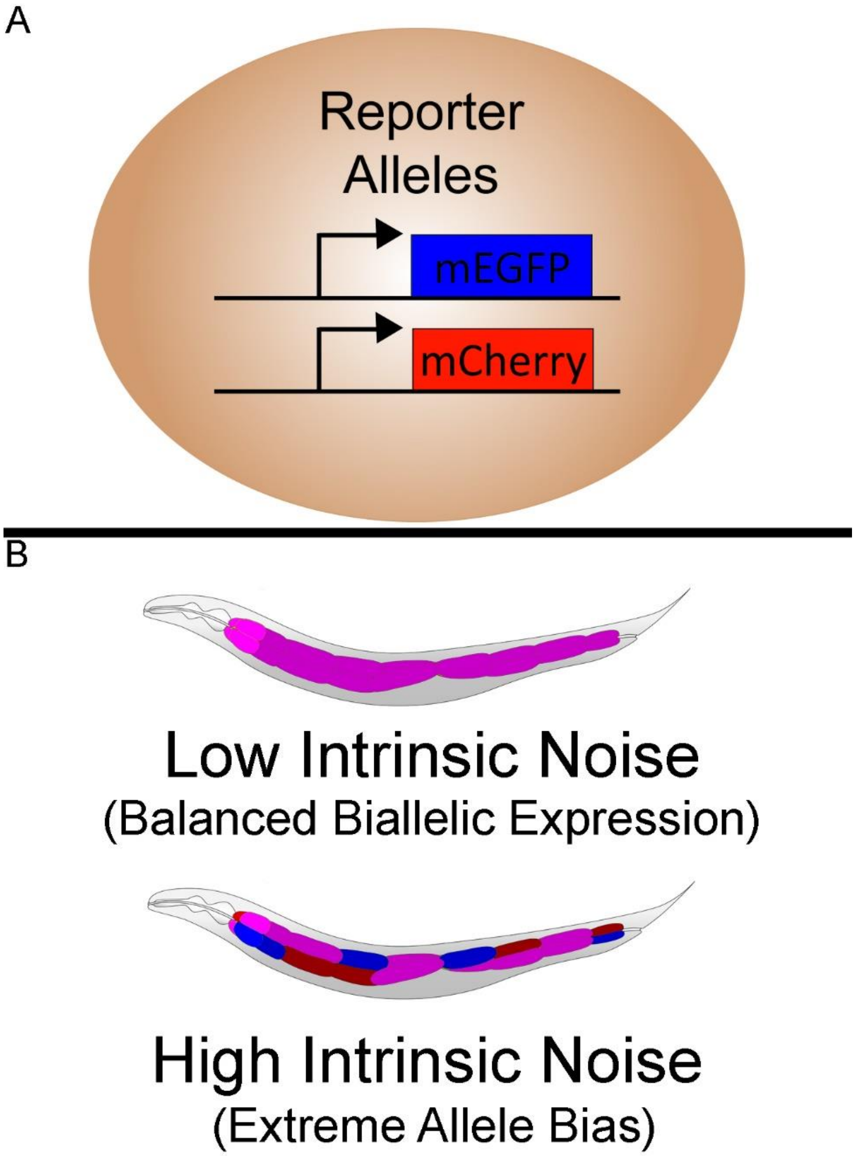


**Fig. S1. Colorblind Corrected Experimental Schematics. A)** shows a cartoon detailing that cells contain two otherwise identical copies of alleles encoding two spectrally distinct fluorescent proteins; here mEGFP is false colored blue to obviate red/green color blindness. **B)** shows cartoon examples of what high and low intrinsic noise would look like among the twenty cells comprising the worm intestine tissue. The top panel shows low intrinsic noise, which appear as purple cells (or somewhat pink), because purple is the balanced blend of red and blue. The lower panel shows high intrinsic noise, which appears as an array of randomly biased and balanced cells; some cells appear purple, and some cells are mostly, or all, red or blue. We will happily recolor any raw microscopic images and send representative or desired images of particular z slices to any interested colorblind individuals.

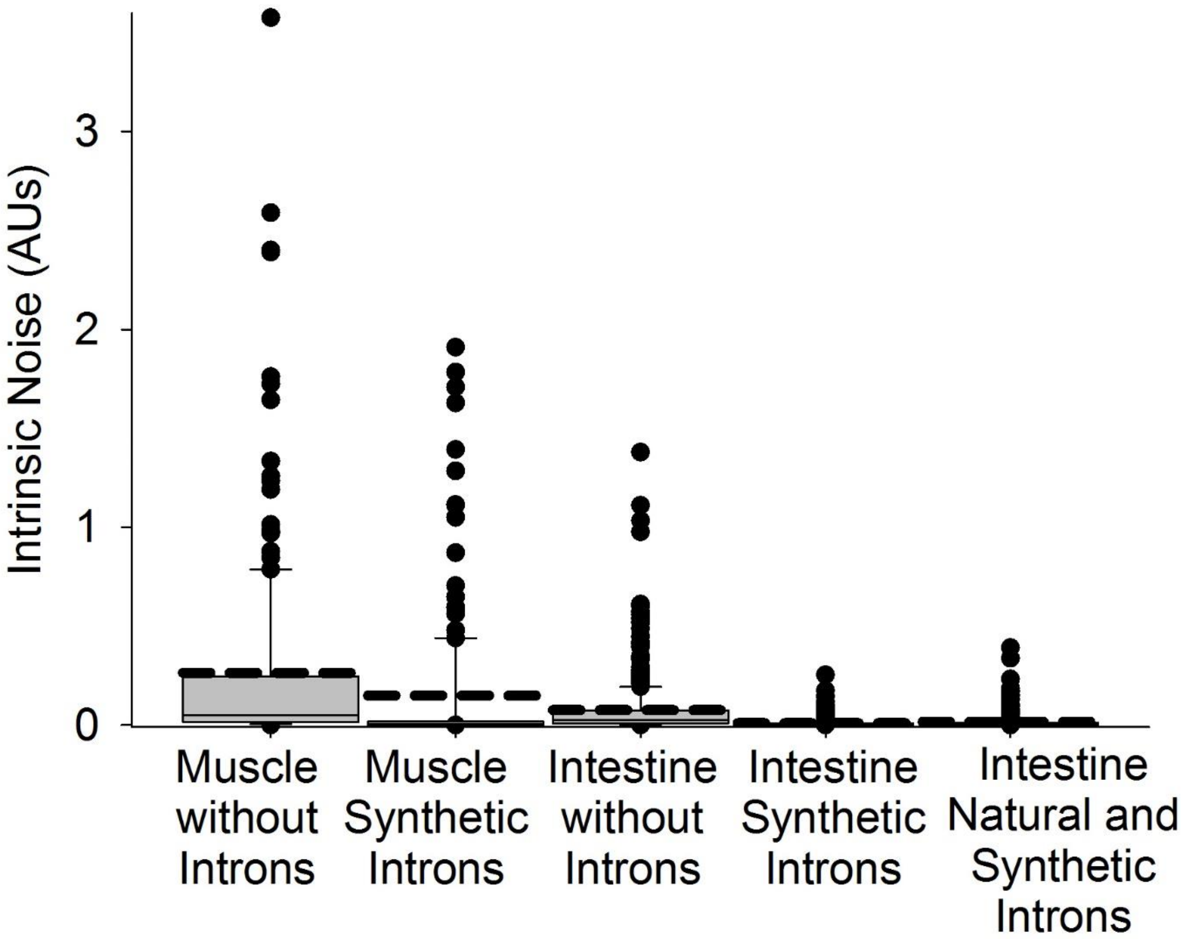


**Fig. S2. Effect of Introns and Cell Types on Intrinsic Noise.** A full scale boxplot showing outliers is displayed. Boxplot shows effects of having introns in alleles and/or different cell types on intrinsic noise.

.

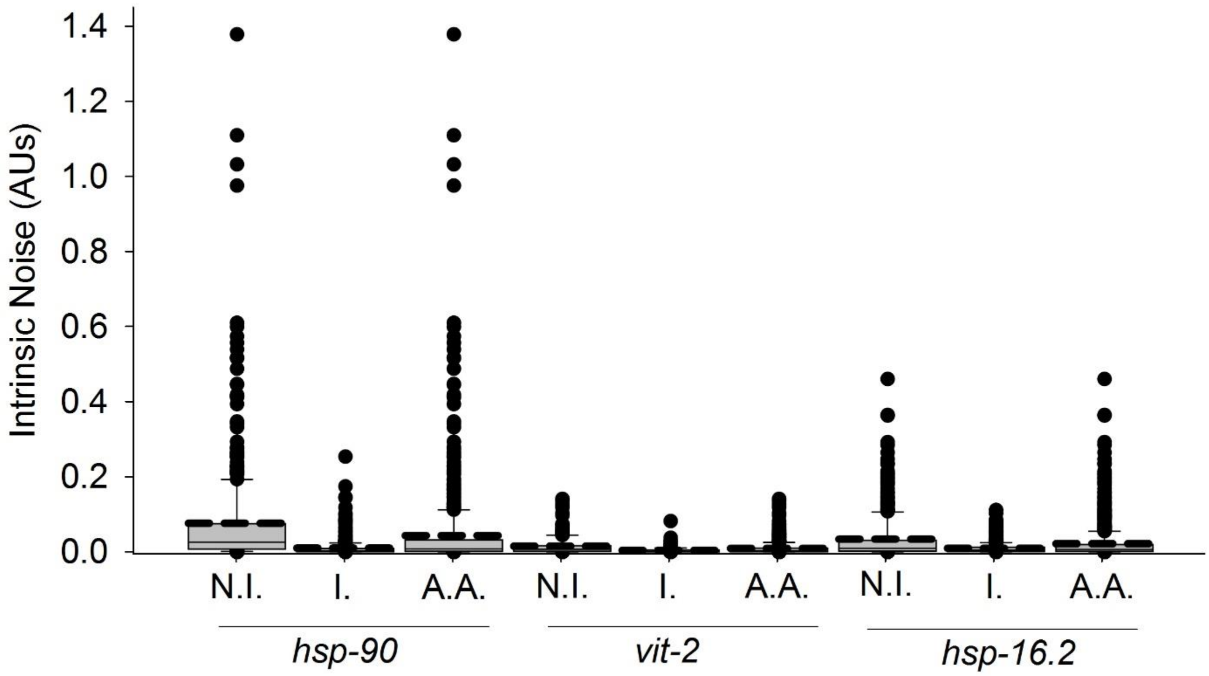


**Fig. S3. Effects of Promoters and Introns on Intrinsic Noise in Intestine Cells.** A full scale boxplot showing outliers is displayed. Boxplot shows effects of having introns in alleles and/or different promoters controlling gene expression on intrinsic noise. N.I. abbreviates “no introns in alleles”, I. abbreviates “introns in alleles”, and A.A. abbreviates “all alleles included”. Promoter controlling gene expression is designated below each trio of groups of alleles for which expression was controlled by that promoter.

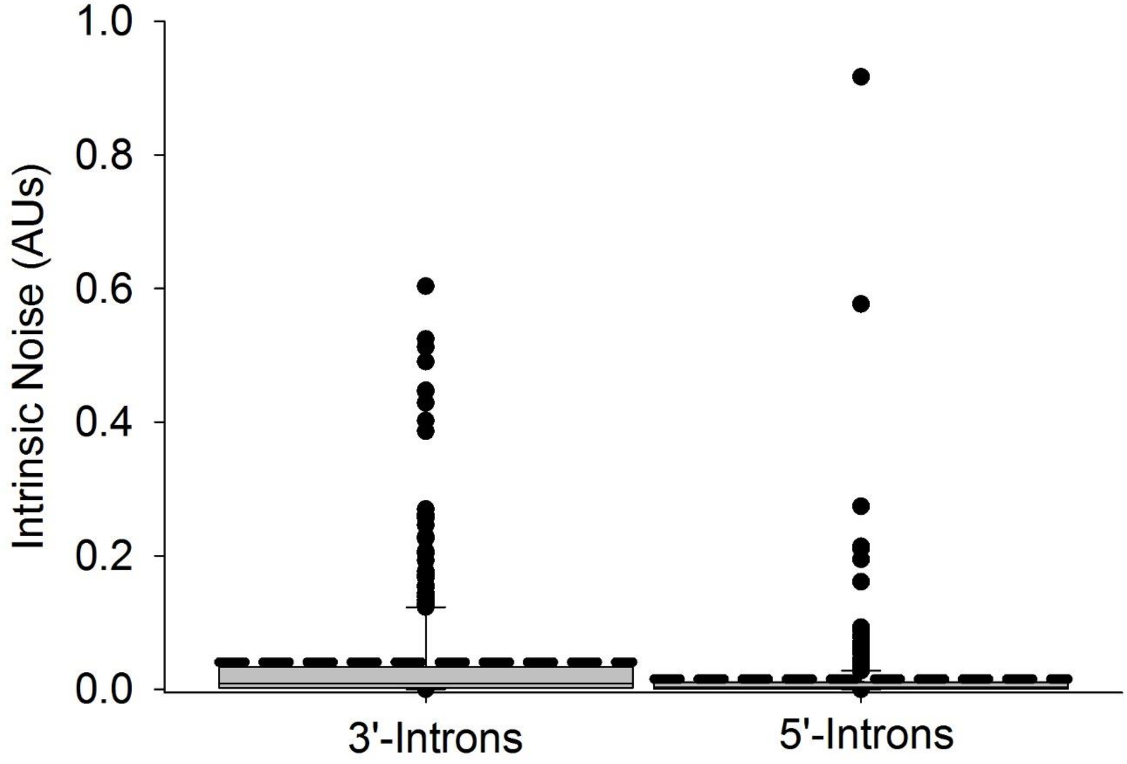


**Fig. S4. Effect of Intron Position on Intrinsic Noise.** A full scale boxplot showing outliers is displayed. Boxplot shows effect of having introns in a relatively 5’ or 3’ position in the coding sequence of fluorescent proteins on intrinsic noise.

| Strain Name | Reporter Allele | Genotype |
| --- | --- | --- |
| ARM136 | *$P_{hsp-90}$::mCherry* | *hutSi2631[$P_{hsp-90}$::mcherry::$T_{unc-54}$]* |
| ARM134 | *$P_{hsp-90}$::mEGFP* | *hutSi2651[$P_{hsp-90}$::mEGFP::$T_{unc-54}$]* |
| ARM133 | *$P_{hsp-90}$::mEGFP* with the three classic synthetic introns | *hutSi2661[$P_{hsp-90}$::mEGFP w/ 3 synthetic introns::$T_{unc-54}$]* |
| ARM210 | *$P_{hsp-90}$::mEGFP* 5' intron | *wamSi210[$P_{hsp-90}$::mEGFP w/ 5'intron::$T_{unc-54}$]* |
| ARM211 | *$P_{hsp-90}$::mEGFP* 3' intron | *wamSi210[$P_{hsp-90}$::mEGFP w/ 3'intron::$T_{unc-54}$]* |
| ARM212 | *$P_{hsp-90}$::mCherry* 5' intron | *wamSi212[$P_{hsp-90}$::mCherry w/ 5'intron::$T_{unc-54}$]* |
| ARM213 | *$P_{hsp-90}$::mCherry* 3' intron | *wamSi213[$P_{hsp-90}$::mcherry w/ 3'intron::$T_{unc-54}$]* |
| ARM137 | *$P_{hsp-90}$::mCherry* natural introns | *wamSi137[$P_{hsp-90}$::mcherry w/3 synthetic introns::$T_{unc-54}$]* |
| ARM135 | *$P_{hsp-90}$::mCherry* synthetic introns | *hutSi2642[$P_{hsp-90}$::mEGFP w/introns::$T_{unc-54}$]* |
| ARM147 | *$P_{vit-2}$::mCherry* | *hutSi2511[$P_{vit-2}$::mCherry::$T_{unc-54}$]* |
| ARM148 | *$P_{vit-2}$::mCherry* 3 introns | *hutSi2581[$P_{vit-2}$::mCherry w/ 3 synthetic introns::$T_{unc-54}$]* |
| ARM145 | *$P_{vit-2}$::mEGFP* | *hutSi2611[$P_{vit-2}$::mEGFP::$T_{unc-54}$]* |
| ARM146 | *$P_{vit-2}$::mEGFP* 3 introns | *hutSi2621[$P_{vit-2}$::mEGFP w/ 3 synthetic introns::$T_{unc-54}$]* |
| ARM139 | *$P_{hsp-16.2}$::mCherry* | *hutSi2552[$P_{hsp-16.2}$::mCherry::$T_{unc-54}$]* |
| ARM140 | *$P_{hsp-16.2}$*::mCherry 3 introns | *hutSi2561[$P_{hsp-16.2}$::mCherry w/ 3 synthetic introns::$T_{unc-54}$]* |
| ARM138 | *$P_{hsp-16.2}$*::mEGFP | *hutSi2591[$P_{hsp-16.2}$::mEGFP w/ 3 synthetic introns::$T_{unc-54}$]* |
| ARM141 | *$P_{hsp-16.2}$*::mEGFP 3 introns | *hutSi2601[$P_{hsp-16.2}$::mEGFP w/ 3 synthetic introns::$T_{unc-54}$]* |

**Table S1.**
Table S1 lists strain names, descriptions and genotypes of reporter gene bearing *C. elegans* strains we created and used in this study.

| Crosses | Description |
|---|---|
| ARM136 x ARM134♂ | *hsp-90* no introns |
| ARM135 x ARM133♂ | *hsp-90* introns |
| ARM212 x ARM210♂ | *hsp-90* 5’ intron |
| ARM213 x ARM211♂ | *hsp-90* 3’ intron |
| ARM137 x ARM133♂ | *hsp-90* promoter; synthetic introns in mEGFP/natural *hsp-90* introns in mCherry |
| ARM147 x ARM145♂ | *vit-2* no introns |
| ARM148 x ARM146♂ | *vit-2* introns |
| ARM139 x ARM138♂ | *hsp-16.2* no introns |
| ARM140 x ARM141♂ | *hsp-16.2* introns |

**Table S2.**
Table S2 lists the initial crosses used to generate stocks of heterozygous animals that were maintained as hermaphrodites through repeated selection of heterozygous hermaphrodites.

**Data S1. (separate file)**

An excel spreadsheet detailing raw, normalized and scaled expression values; the sheets also include calculations for intrinsic noise and intrinsic noise strength. Allele expression levels from individual intestine or muscle cells are listed in different tabs, with each tab detailing the genotype and cell type for muscle cells; all other cells are intestine cells from rings 1-4. Muscle cells were sampled from muscle rows adjacent to intestine rings 1-4.

**Data S2. (separate file)**

An excel spreadsheet listing intronless, protein coding genes in modern drafts of the *C. elegans* and human genomes in different tabs. Additional spreadsheets in separate tabs detail molecular function and biological process GO terms enrichment for intronless human and worm genes. Another spreadsheet in another separate tab details genes found to have chromatin signature or RNA-seq based evidence of monoallelic expression, listed at the database for monoallelic gene expression (dbMAE).

### Materials and Methods

*Molecular Cloning*

We generated all of the DNA constructs by 3-fragment DNA assembly in yeast, using a protocol that we recently published [1]. Briefly, we mixed competent yeast with promoter and GFP or mCherry PCR products and 60ng of linearized expression vector BSP188. This expression vector contains the unc-54 terminator and chromosome II MosSCI homology arms for integration at ttTi5605. For promoters, we used worm gDNA to amplify sequence 2Kb upstream of the ATG for *hsp-90*, 392bp upstream of the ATG for *hsp-16.2*, and 4Kb upstream of the ATG for *vit-2*. Intronless transgenes were assembled by overlap extension PCR using intron-containing transgenes as template. We rescued assembled DNA into *E. coli* and sequence verified the final assembled plasmids.

*Genome Editing/Strain Creation*

Plasmid DNAs were then used in microinjection-based MosSCI transgenesis to insert the reporter cassettes into the genome at the ttTi5605 locus on Chromosome II in strain RBW6699, which is an outcrossed version of EG6699 (@ 50ng/µL of repair template + coinjection and selection markers [2-4]). Thus, all reporter allele cassettes were integrated into the same autosomal ttTi5605 Chromosome II locus, including the relatively 3' *C. briggsae unc-119* phenotypic rescue marker. We generated some strains *de novo* for this study and used other, more outcrossed versions of strains we made and first reported in [5], due to the novel nature of our results. Thus, we outcrossed each strain reported here with N2 wild-type animals a minimum of three times. The resulting strain names and genomic insertion designations are shown in Supplemental Table 1.

*Animal Husbandry*

Animals were cultured as previously described [6]. Briefly, we maintained all strains in 10cm petri dishes on NGM seeded with OP50 *E. coli* in an incubator at 20°. All strains used in this study are listed in Table S1. A table of crosses can be found in Table S2. To begin the crosses, we heat shocked 10cm plates containing 20 L4 animals per plate at 30 degrees for 5-6 hours to produce males. We mixed young males with L4 hermaphrodites on 3cm mating plates at a ratio of 3 to 5 males per hermaphrodite. 24 hours post-mating, individual P0 hermaphrodites were picked onto fresh 6cm NGM plates to lay progeny. L4s of the F1 generation were picked onto 10cm plates to lay a mix of heterozygous and homozygous progeny. We screened for heterozygous animals that express both GFP and mCherry on a fluorescence stereoscope. We maintained heterozygotes by picking them away from homozygous animals and onto fresh, OP50-seeded NGM growth plates each generation. We performed experiments on heterozygous animals that were at least three generations beyond the initial cross to avoid paternal allele expression bias. To synchronize animals for experiments, we conducted 2 hour egg lays onto 10cm NGM plates (10 heterozygous animals per plate).

*Microscopy*

We washed day 2 adult animals into M9 media with tricane/tetramisole [7] and loaded animals into 80-lane microfluidic devices that we recently described [1]. These devices immobilize worms in 80 separate channels. We imaged only those animals that randomly immobilized with their left side facing the cover slip, to which the fluidic device was bonded, which put intestine cells in rings I through IV closest to the microscope objective. The muscle cells were on the oblique, dorsal and ventral sides of the animals, and less easy to observe in the lateral orientation that animals tend to assume on slides and in these devices.

We imaged animals as we previously described [7]. Briefly, to image the animals, we used a 40X 1.2 NA water objective on a Zeiss LSM780 confocal microscope. We excited the sample with 488 and 561 nm lasers and collected light from 490-550nm for mEGFP signal and from 580-640nm for mCherry signal. We focused on the same field of view for each animal- starting from the posterior of the pharynx to the first half of cells in intestinal ring IV. We collected images of the entire z depth of each animal, from one side to the other, using two micrometer step size and a two micrometer optical slice as we have previously described [7].

*Image Cytometry*

Our image cytometry was conducted in a manner that we have previously described in detail in [7]. Our image cytometry consists of manual cell identification and annotation, with a semiautomatic quantification step. Briefly, we first determined the orientation of the animals in images and then identified individual intestine or muscle cells. We then measured signal within an equatorial slice of the cell's nucleus, as a proxy for the whole cell. Nuclear signal of freely diffusing monomeric fluorescent protein is nearly perfectly correlated with the cytoplasmic contents [7]. We used the ImageJ software as well as custom built Nuclear Quantification Support Plugin for nucleus segmentation and signal quantification. We call the ImageJ plugin *C. Entmoot* (Alexander Seewald, Seewald Solutions, Inc., Vienna), as the program segments as the result of a meeting of decision trees, much like an Ent Moot in J. R. R. Tolkien's Lord of the Rings (a meeting of fantasy tree-like humanoid beings to make a collective decision).

*Data Processing and Noise Calculations.*

Here we measured intrinsic noise by measuring the expression level of differently colored reporter alleles in two-day old adult animals that appear to be in a steady-state of gene expression. Intrinsic noise is essentially the quantitative measure of relative deviation from the diagonal trend line. Intrinsic noise measures how deviant a pair of reporter alleles from the average ratio among groups of cells, thus quantifying how probable it is to observe biased or monoalleleic expression for a given gene (pair of alleles) in a given population of cells (e.g, muscle cells or intestine cells). The assumptions of our intrinsic noise model are the same as the assumptions in [7]. We sometimes used 8-bit or 16-bit file settings during data collection. We normalized expression level data for each allele to per-experiment means and scaled the data by an arbitrary value of 100. We calculated intrinsic noise and intrinsic noise strength as detailed in [8-10]. Data with calculations are available as Supplementary Data File, Data S1, in Excel format.

We conducted three independent experiments for muscle cells and four independent experiments for all other datasets. We collected data from eight of the exact same intestine cells in intestine rings I-IV (avoiding the distal cells in ring I), and sometime included additional intestine cells in ring V. For experiments measuring muscles, we collected data from six intestine-adjacent muscle cells, from either, or both, the dorsal left muscle row, and the ventral left muscle row. In each experiment we collected data from intestine or muscle cells from at least ten different animals per group. We manually curated data from thousands of cells, with a minimum of 180 cells (for muscles) and as many as 398 cells in total for each experimental groups. See also statistical analyses.

*Plotting and Statistics*

We used SigmaPlot 12.5 (Systat Software, Inc., San Jose) for all plotting and statistical analyses of noise or noise strength. Briefly, for experiments with multiple groups, we ran ANOVA followed by appropriate parametric or nonparametric post hoc tests, detailed below. For experiments with only two groups, we ran a nonparametric Mann Whitney U test.
